# Maternal body condition affects the response of larval spined toads’ faecal microbiome to a widespread contaminant

**DOI:** 10.1101/2023.12.18.572122

**Authors:** Sabrina Tartu, Nicolas Pollet, Isabelle Clavereau, Gauthier Bouchard, François Brischoux

## Abstract

Glyphosate’s primary metabolite, aminomethylphosphonic acid (AMPA), is the most detected pollutant in surface waters. Recent studies have raised concerns about its toxicity, yet underlying mechanisms remain poorly understood. A disruption of the gut microbiome, which plays a crucial role in host health, could mediate most of the adverse effects. We investigated the impact of AMPA exposure on the gut microbiome of spined toad tadpoles (*Bufo spinosus*). We hypothesized that AMPA could alter the gut microbiota composition and that these effects could depend on the microbiota source. We exposed tadpoles to minute concentrations of AMPA and analyzed their faecal microbiota using 16S rRNA gene sequencing as a proxy of the gut microbiota. AMPA exposure decreased the gut bacterial biomass and affected the bacterial community composition of tadpole’s faeces. Furthermore, we observed interactions between AMPA exposure and maternal body condition on the Bacteroidota and Actinobacteriota phyla abundances. This suggests a maternal effect on early-life microbial colonizers that could influence the response of the gut microbiome to AMPA. These findings highlight the importance of considering the gut microbiome when studying the effects of environmental contaminants. Further research is needed to elucidate the long-term implications of this microbiome alteration for amphibian health.

## INTRODUCTION

Over the years, regulatory agencies have banned most highly toxic and persistent pesticides, such as the notorious DDT, and replaced them with other, fast-degrading and more species-specific compounds. However, several current-used pesticides and their transformation products still pose ecotoxicological issues (Gonçalves et al., 2021). Aminomethylphosphonic acid (AMPA, CAS No. 1066-51-9) is one of those transformation products that may pose higher risks than its parent compounds (Grandcoin et al., 2017). Although AMPA has two primary sources, phosphonate and glyphosate degradation, through the lysis of the C-P bond and action of the enzyme glyphosate oxidoreductase, respectively (Jaworska et al., 2002; Zhan et al., 2018), its origin in surface water and groundwater is mainly linked to the latter (Carles et al., 2019; Struger et al., 2015). Importantly, AMPA is detected much more frequently (20-50% more detected) and is more persistent in the environment (half-life 2-8 times longer) than glyphosate, with concentrations in surface water generally ranging between 0.2 and 5 µg L^-1^ (Duke, 2020; Grandcoin et al., 2017; Kolpin et al., 2006; Maggi et al., 2020; Ojelade et al., 2022).

In aquatic organisms, the effects of AMPA exposure are controversial, ranging from low toxicity to numerous adverse effects, including increased mortality, development delay, increased morphological abnormalities, genotoxicity, increased oxidative stress, cardiac defects, hepatic inflammatory response or changes in metabolic activity (Antunes et al., 2017; Barreto et al., 2023; Cheron et al., 2022; Cheron and Brischoux, 2023, 2020; Guilherme et al., 2014; Ivantsova et al., 2022; Levine et al., 2015; Martins-Gomes et al., 2022; Matozzo et al., 2018; Tartu et al., 2022; Zhang et al., 2021). However, despite the numerous adverse effects reported on non-target organisms, AMPA is not under particular scrutiny, and the underlying mechanisms involved in its ecotoxicity are poorly identified.

The gut microbiome is a crucial endpoint for ecotoxicological studies (Claus et al., 2016; Evariste et al., 2019). The trillions of microorganisms that have colonized a host compose its microbiota, with more than 90% of this commensal, mutualistic and symbiotic microbe community located in the hosts’ gut (Sharpton, 2018). The gut microbiota composition depends on both horizontal (e.g. habitat, diet, conspecifics) and vertical transmission (e.g. parents) in vertebrates (Comizzoli et al., 2021; Moeller et al., 2018; Murphy et al., 2023; Robinson et al., 2019; Scalvenzi et al., 2020). A large number of studies have underlined the functional importance of the gut microbiota composition and structure in food digestion, nutrient synthesis, host’s physiology, development, behaviour, or immune system performance (Clemente et al., 2012; Grond et al., 2018; Jiménez and Sommer, 2017).A dysbiosis, consisting of a modification in the composition and function of the gut microbiota in response to a stressor, can alter intestinal permeability, affect physiological performances and immune response, increasing disease susceptibility (Gomaa, 2020; Grond et al., 2018; Warne et al., 2019; Xiong et al., 2019), which could lead to unforeseen hazardous consequences for wild populations.

Because of its mode of action which is to inhibit the enzyme 5-enolpyruvyl-shikimate-3-phosphate synthase (EPSPS) of the shikimate pathway, a metabolic pathway specific to plants but also microorganisms (Herrmann and Weaver, 1999), several studies have tested the effects of glyphosate exposure on vertebrates and invertebrates gut microbiota composition (Blot et al., 2019; Cuzziol Boccioni et al., 2023; Ding et al., 2021; Fréville et al., 2022; Iori et al., 2020; Lehman et al., 2023; Lozano et al., 2018; Mesnage et al., 2021; Motta et al., 2018; Owagboriaye et al., 2021; Puigbò et al., 2022; Ruuskanen et al., 2020; Walsh et al., 2023). Yet, only two have focused on the effects of AMPA on invertebrate models and reported slight effects on gut microbiota (Blot et al., 2019; Iori et al., 2020). Considering the widespread presence of AMPA in the environment and its ability to affect microorganism growth by inhibiting bacterial cell wall biosynthesis (Atherton et al., 1982; Azam and Jayaram, 2016; Carles et al., 2019; Coupe et al., 2012; Poiger et al., 2017), there is still a lack of research linking the harmful effects of AMPA to a gut microbiome dysbiosis.

Amphibians face a higher overall extinction rate than other vertebrates (Harfoot et al., 2021; Hoffmann et al., 2010). Intensive agriculture is a significant threat to which they are exposed, contributing to habitat loss and pollution (Harfoot et al., 2021; Rollins-Smith, 2020; Stuart et al., 2004; Wake and Vredenburg, 2008). Their reproductive migrations, water-dependent breeding, aquatic larval stages, highly permeable skin and limited movements make amphibians particularly vulnerable to environmental changes and highly relevant models to study the effects of environmental contamination (Langlois, 2021; Stebbins and Cohen, 1995).

Previous studies conducted on the spined toad, *Bufo spinosus*, have associated AMPA exposure with numerous adverse effects, such as higher embryonic and larval mortality, increased oxidative stress in tadpoles (Cheron et al., 2022; Cheron and Brischoux, 2023, 2020) and altered colouration in adult males (Tartu et al., 2023). In addition, in a companion study using the same individuals as the present one, we reported that environmentally relevant concentrations of AMPA (0.4 µg L^-1^) led to increased deformity rate upon hatching and increased development length, with effects depending on the habitat of origin of the parents (agricultural *versus* forest, Tartu et al., 2022). The composition of gut microbiota is tightly associated with growth rate and metamorphosis in anuran species (Emerson and Woodley, 2024; Lv et al., 2023; Park et al., 2023). Therefore, we tested whether the previously observed adverse effects of AMPA on spined toad tadpoles could originate from gut microbiota dysbiosis. Moreover, we further hypothesized that AMPA-mediated effects on the gut microbiota of tadpoles would be dependent on parent’s body-condition (as a proxy of parental diet and quality) and habitat of origin (agricultural or forest).

## MATERIAL AND METHODS

### Fieldwork

The 120 spined toad tadpoles used presently are a subset of the 240 individuals employed in a previous companion study (Tartu et al., 2022). Study sites, parent captures, and housing conditions have been described previously (Tartu et al., 2022). Briefly, between 28/01/2021 and 22/02/2021, we searched for spined toad amplectant pairs at night in two agricultural sites and in two sites surrounded by woodlands (see supporting information in (Tartu et al., 2022)). We collected the ten first spotted couples to guarantee comparable inter-site individual quality. We then returned breeding pairs to the laboratory until females lay their eggs (**Figure S1**). After oviposition, we measured the snout-vent length of both parents with a calliper and weighed them on a precision scale. We calculated the scaled mass index (SMI) developed by Peig and Green (2009) to assess parental body condition as SMI accurately reflects amphibian energy stores in anuran species (Băncilă et al., 2010; Landler et al., 2023; MacCracken and Stebbings, 2012; Zhelev and Minchev, 2023). We described how we categorized males and females into the thin and fat groups in the “statistical analyses” section below. Finally, we released the pairs in their breeding site after body measurements.

### Housing conditions and AMPA treatment

We obtained six segments of 34 eggs for each clutch and placed each segment in individual aquariums with 2L of dechlorinated tap water. Among the six segments, we exposed three to 0.4 µg L^-1^ (± 0.01 µg L^-1^) of AMPA (AMPA group), and we kept the remaining three in dechlorinated tap water as a control group. We obtained the AMPA solution by dissolving commercial crystalline powder (Aminomethylphosphonic acid, 99% purity, ACROS ORGANICS™) with dechlorinated tap water. An independent accredited analytical laboratory confirmed AMPA concentration in water (QUALYSE, Champdeniers-Saint-Denis, France). As evidenced by data available from French national surveys conducted between 2020 and 2023 on 380 samples obtained from 67 rivers of the Deux-Sèvres Region, AMPA concentrations ranged from 0.02 to 3.4 µg L^-1^ (Naïades, 2023), the selected AMPA concentration (0.4 µg L^-1^) in the present study represents actual concentrations found in surface water in our study area.

Upon hatching, we randomly isolated one tadpole and released the remaining tadpoles in their parents’ breeding pond. We housed each selected tadpole individually (n=240 in total; see **Figure 1, S1**) in a 2L aquarium with either dechlorinated tap water or AMPA according to the treatment experienced during embryonic development. Consequently, tadpoles were exposed to the same treatment during embryonic and larval development. We checked tadpoles daily and monitored their development through six key Gosner stages (25, 30, 37, 41, 42 and 46 (Gosner, 1960)). The effects of AMPA exposure on tadpole development according to the parent’s origin (forest vs agricultural) have been published previously (Tartu et al., 2022).

**Figure 1:**
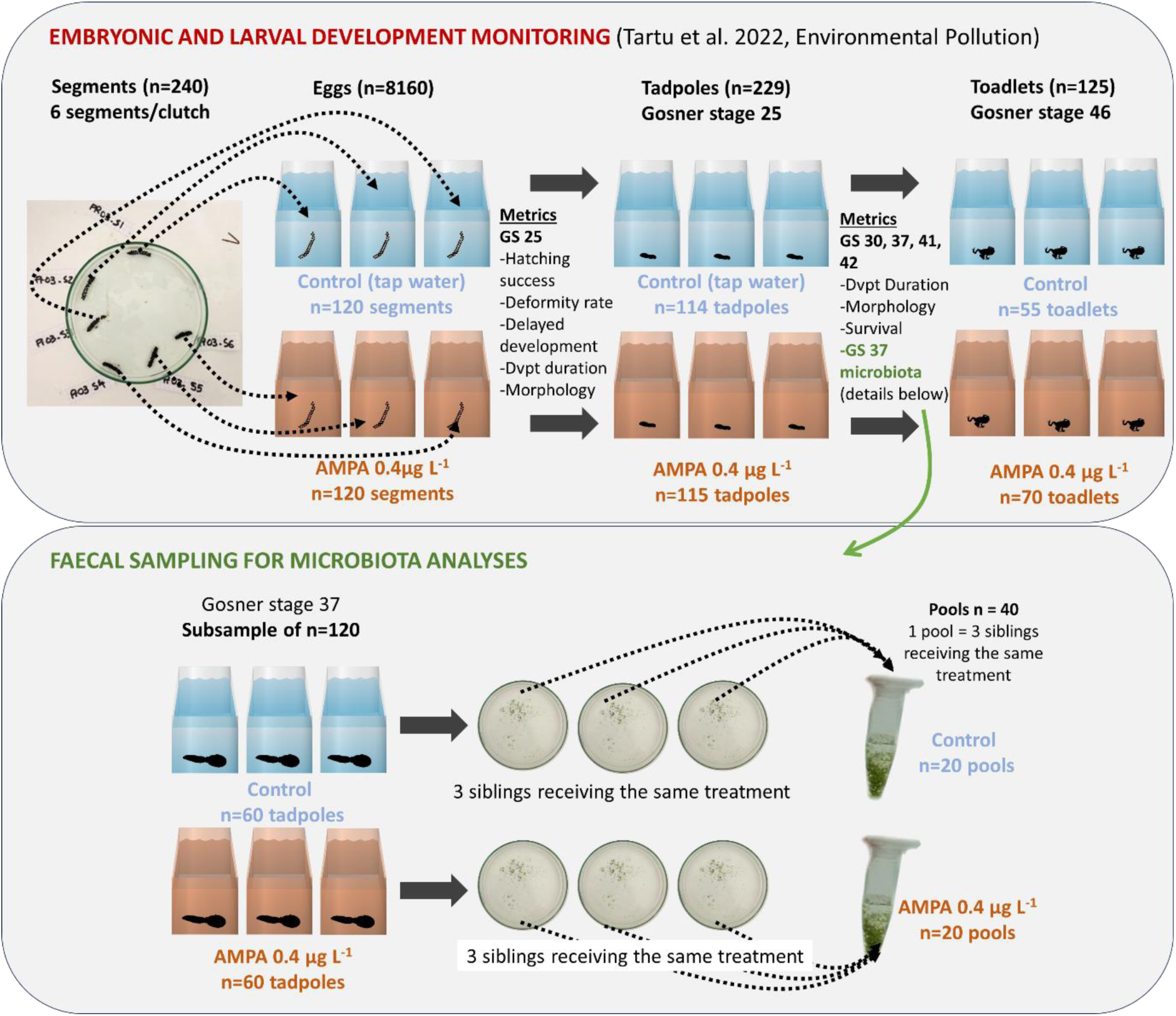
Common-garden experiment conducted on spined toad *Bufo spinosus* tadpoles. The upper panel shows our experimental design and the metrics monitored for each Gosner stage (GS). From each clutch, we exposed three segments to control conditions (dechlorinated tap water) and three segments to AMPA (0.4 µg L^-1^). From each segment that produced at least one viable tadpole, one was kept and monitored until metamorphosis (Gosner stage 46). Lower panel: on a sub-sample of tadpoles (n=120), we collected the faeces of individuals when they reached GS 37. We then pooled the faeces of three siblings who had received the same treatment in a single tube (n=20 control, n=20 AMPA) to conduct microbiota analyses.

From the egg stage until metamorphosis, the tadpoles were kept under simulated 12:12 h day and night in a room at 17°C to avoid any basal metabolism and development variation. Water was changed weekly and 0.4 µg L^-1^ AMPA was added to the AMPA group tanks after each water change. Upon hatching, we fed tadpoles with organic ground spinach *ad libitum*. Ethics committees approved this study (permits APAFIS#13477–2018032614077834 and DREAL/2020D/8041).

### Faecal sampling for microbiome analyses

Although there are some controversies about using faecal microbiome to reflect gut microbiome comprehensively (Tang et al., 2020), we still privileged this non-invasive sampling method to release the toadlets upon metamorphosis. During pre-metamorphic larval development, spined toad tadpoles are the most active swimming and feeding at Gosner stage 37 (Cheron et al., 2021), resulting in a more significant production of faeces. Therefore, we started sampling tadpoles’ faeces for gut microbiota analyses once they reached Gosner stage 37. To do so, we collected faeces accumulated in the bottom of the aquarium since the last water change (4-6 days) using sterile pipettes for half of the individuals (n = 120). We placed faeces in a sterile petri dish, then transferred them by pipetting with filter tips to a sterile microtube and added twice their volume of DNA/RNA Shield™ (Zymo Research). To increase the quantity of faeces per sample and the genetic diversity of microbial communities, we pooled the faeces from three sibling tadpoles receiving the same treatment (control or AMPA) in one tube. We thus obtained 40 pools (**Figure 1**). DNA/RNA Shield™ preserves the nucleic acids integrity of samples at ambient temperatures. We therefore kept the pooled faecal samples at room temperature until analyses.

### DNA extraction, libraries and sequencing

We extracted DNA using the ZymoBIOMICS DNA/RNA Miniprep kits according to the manufacturer’s instructions. We added 1 µL of the Zymobiomics Spike-in control I (High microbial Load D6320) to each sample as *in situ* positive control (Galla et al., 2023) before lysis using a Precellys Evolution Touch equipped with a cryolys module to perform six steps of bead beating at 10,000 rpm during 10 s at 0°C, with 30 s pause between each step. We included a set of samples from artificial communities as positive controls (Zymobiomics Microbial community standard (D6300) and log distribution (D6310)) and negative controls (tap water used to rear tadpoles and nuclease-free water). We controlled the quality and quantity of extracted DNA (3180 ± 2479 ng DNA per faeces sample) using spectrophotometry (Nanodrop) and fluorometry (Qubit). We amplified the V1-V8 portion of the 16S rDNA gene by PCR using tailed primers tBACT27F (5’-TTTCTGTTGGTGCTGATATTGCAGAGTTTGATCCTGGCTCAG-3’) and tBACT1391R (5’-ACTTGCCTGTCGCTCTATCTTCGACGGGCGGTGWGTRCA-3’). We ran PCR in triplicate reactions of 25 µL assembled under a PCR laminar flow cabinet using the following conditions: 40 ng DNA, 0.3µM each primer, 0.5 mM dNTPs, 1 Unit of tiAmplus DNA Polymerase HotStart (Roboklon), 1X tiAmplus Buffer containing 25 mM MgSO_4_. The PCR program was 93°C for 2 min, followed by ten cycles of 10s at 93°C, 30s at 57°C, 120s at 68°C and 25 cycles of 10 s at 93°C, 30s at 65°C, 120s at 68°C and 7 min at 68°C. We monitored amplification products using regular gel electrophoresis before pooling the triplicate PCR and purification using magnetic beads (Macherey-Nagel NucleoMag cleanup). After the first round of PCR (16S), we obtained a mean quantity of 1.82 ± 0.59 µg of pooled PCR products after purification. We used 100 ng of each PCR product for a second round of PCR (multiplexing) and finally obtained 3.4 ± 1.3 µg of purified product. We then used the PCR barcoding expansion pack (EXP-PBC096, Oxford Nanopore Technology) and the ligation sequencing kit (SQK-LSK109) to prepare a sequencing library of which 150 ng were loaded on the flow cell (R9.1) and sequenced using a MinIon device (Oxford Nanopore Technology). Two libraries were sequenced on two different flowcells, yielding 5,043,247 and 5,184,836 usable reads after demultiplexing. The sequenced datasets generated and analyzed during the current study are available in the EMBL Nucleotide Sequence Database (ENA) at https://www.ebi.ac.uk/ena/data/view/PRJEB71117.

### 16s rDNA sequence analysis

We mapped individual reads using minimap2 on the SILVA SSU database version 138. We retained only reads mapping with a coverage of 75% and an alignment length between 1000 and 2000 nt to a SILVA entry identified by at least two reads in a given sample. We ran abundance analyses using OTUs identified by a prevalence of at least four with a minimum read depth of three.

### Analysis of spike control

We analyzed four samples as negative controls to monitor the inputs from the dechlorinated tap water used to raise tadpoles (two samples) or any contaminants arising from sample manipulation, DNA extraction, PCR amplification and sequencing library construction (two samples). As in all samples, we included spiked bacteria (D6320, ZymoBIOMICS) in the controls we analyzed in two independent sequencing runs. We obtained 19,270 and 28,848 reads in the water samples and 26,579 and 38,633 reads in the kit samples, from which respectively 98.25; 98.73; 99.77; and 99.71% were identified as the spiked bacteria *Imtechella* and *Allobacillus*. Therefore, the bacterial inputs from the dechlorinated tap water to faecal samples are negligible; spiked bacteria only represented between 0.002-0.06% in faecal samples. The maximum number of reads from another OTU was between 11 and 140, and we found at least one read attributed to 25-78 phylogenetically unrelated OTUs. The ratios of *Allobacillus* to *Imtechella* reads were 0.84-1.05, compared with the theoretical expectation of 2.3.

In conclusion, as expected, we found that a vast majority of the reads from these spike controls identified the two bacteria from the spike. Yet, we observed a trace level of reads derived from phylogenetically unrelated OTUs that artificially increased the number of observed OTUs. We observed that two samples out of seven expected to be devoid of spike-in contained one and three reads from *Allobacillus* but none from *Imtechella*. The sample containing one read may represent a case of spillover since it is located next to a D6320 sample. We computed a spike in OTU abundance from the mean of 100 rarefactions to account for sample read depth variations. We also analyzed the spike-in fraction (D6320) out of the total bacterial abundance to estimate bacterial biomass *in situ* (Jones et al., 2015; Stämmler et al., 2016; Tkacz et al., 2018).

### Statistical analyses

First, we obtained an estimation of the ratios of absolute endogenous bacteria by analyzing the abundance of spike-in controls according to the treatment (AMPA vs control) and their interaction with the parental capture site (agricultural vs forest) and parental (maternal and paternal) body condition (Hornung et al., 2019; Stämmler et al., 2016). We used linear mixed effect models (LME) with clutch identity as a random factor. Then, we evaluated the similarity of bacterial communities between the different treatments and their interaction with the parental capture site and parental body condition with unweighted (based on the presence or absence of observed bacterial taxa) and weighted (based on the abundance of observed bacterial taxa) UniFrac distances (Lozupone et al., 2006). We constructed discrete body condition classes to compare spike-in control abundances and bacterial communities according to treatment while considering parent body condition. To do so, we used an objective categorization method by grouping all the individuals with an SMI ≤ median SMI value as “thin” and all individuals with an SMI > median SMI value as “fat”. We used the non-parametric Kruskal-Wallis (KW) test when comparing UniFrac distances according to more than two variables. Post-hoc analyses were performed using Dunn’s test with the Bonferroni adjustment method. We then performed principal coordinate analysis (PCoA) based on unweighted and weighted UniFrac distances using the ‘*Phyloseq*’ and ‘*ade4*’ packages (Dray and Dufour, 2007; McMurdie and Holmes, 2012). Second, we extracted the abundances of the nine most represented phyla and inserted them into linear mixed effect models (LME) with clutch identity as a random factor. We used LME to specifically analyze which phyla were influenced by the treatment and potential interaction with other variables (e.g. parentalhabitat and body condition). At last, we conducted Linear discriminant analysis Effect Size (LEfSe) analyses within the phyla and variables previously identified as sensitive to AMPA exposure to determine which microbiome biomarkers characterized the observed differences (Segata et al., 2011) by using the ‘*microbiomeMarker*’ package (Cao et al., 2022). We performed all analyses with R v.4.3.2 (R Core Team, 2019).

## RESULTS

### Faecal microbiota composition of spined toad tadpoles

We identified 664 Operational Taxonomic Units (OTUs) within the 40 faecal samples analyzed. Proteobacteria (537 OTUs) was the most abundant phylum, followed by Bacteroidota (73 OTUs), Actinobacteriota (18 OTUs), Firmicutes (13 OTUs), Verrumicrobiota (7 OTUs), Acidobacteriota (7 OTUs), Campilobacterota (4 OTUs), Desulfobacterota (3 OTUs), Planctomycetota (2 OTUs).

### Treatment effects and their interactions with habitat and parental body condition

We observed a higher abundance of spiked-in bacteria in faecal samples from AMPA-exposed tadpoles as compared to controls (LME, estimate: 20.93 ± 6.39, p=0 .004, **Figure 2**), which indicates an endogenous lower biomass in faecal samples of AMPA treated tadpoles. The interaction of treatment with parental habitat and body condition was unrelated to spike-in control abundances (LME, p>0.560 for all tests). Although AMPA treatment did not overall affect faecal microbiota community structure (unweighted UniFrac distances, Kruskal-Wallis χ^2^= 1.99, df = 2, p = 0.370), it affected its composition (weighted UniFrac distances (χ^2^= 11.0, df = 2, p=0.004, post-hoc test AMPA vs Control: p=0.003, **Figure S2**).

**Figure 2:**
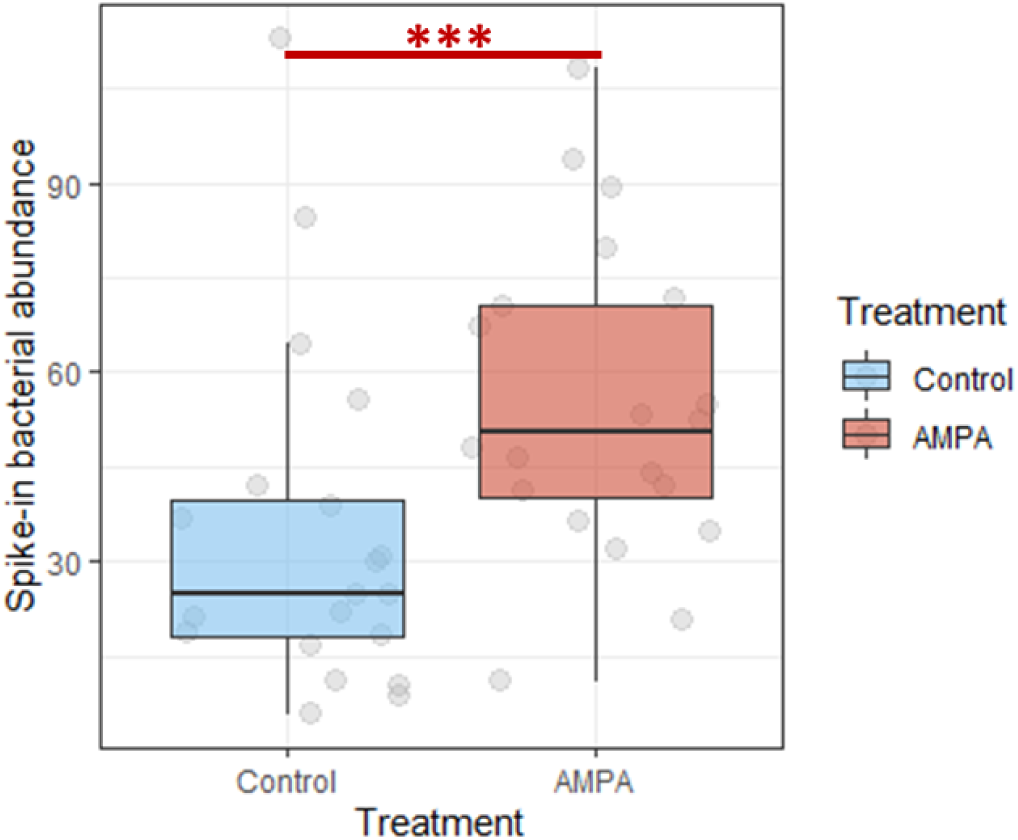
Exogenous spike-in bacterial relative abundance in tadpoles’ faecal samples. The greater abundance of exogenous spiked-in bacteria in samples from AMPA-treated tadpoles reveals an overall lower endogenous bacterial biomass than in the control group. Bacterial abundances were computed as the mean value from 100 rarefactions. Significant differences are represented by *** (p=0.004).

The effect of the treatment on the faecal microbiota community was not exacerbated when consideringthe habitat of the parents (unweighted UniFrac, χ^2^= 11.2, df = 9, p = 0.262; weighted UniFrac, χ^2^= 15.4, df = 9, p = 0.080). Although close to statistical significance for weighted UniFrac distances, pairwise post-hoc comparisons did not reveal any trend (p>0.21 for all tests).

Yet, the AMPA effect depended on maternal body condition (**Figure 3**). More specifically, the structure and composition of faecal microbial communities were affected by AMPA in tadpoles produced by females in better condition. In comparison, this effect was not found in tadpoles produced by females characterized by lower condition(unweighted UniFrac distances, χ^2^= 46.2, df = 9, p = 5.46×10^-07^, post-hoc test: AMPA vs Control in fatter females: p = 0.006, thinner females: p = 0.198; weighted UniFrac, χ^2^=34.1, df = 9, p = 8.53×10^-05^, post-hoc test: AMPA vs Control in fatter females: p = 0.0004, thinner females: p = 0.999, **Figure 3A-D**). Paternal body condition alone or in interaction with the treatment did not significantly influence the tadpole microbiome (p>0.05 for all tests, **Figure 4**).

**Figure 3:**
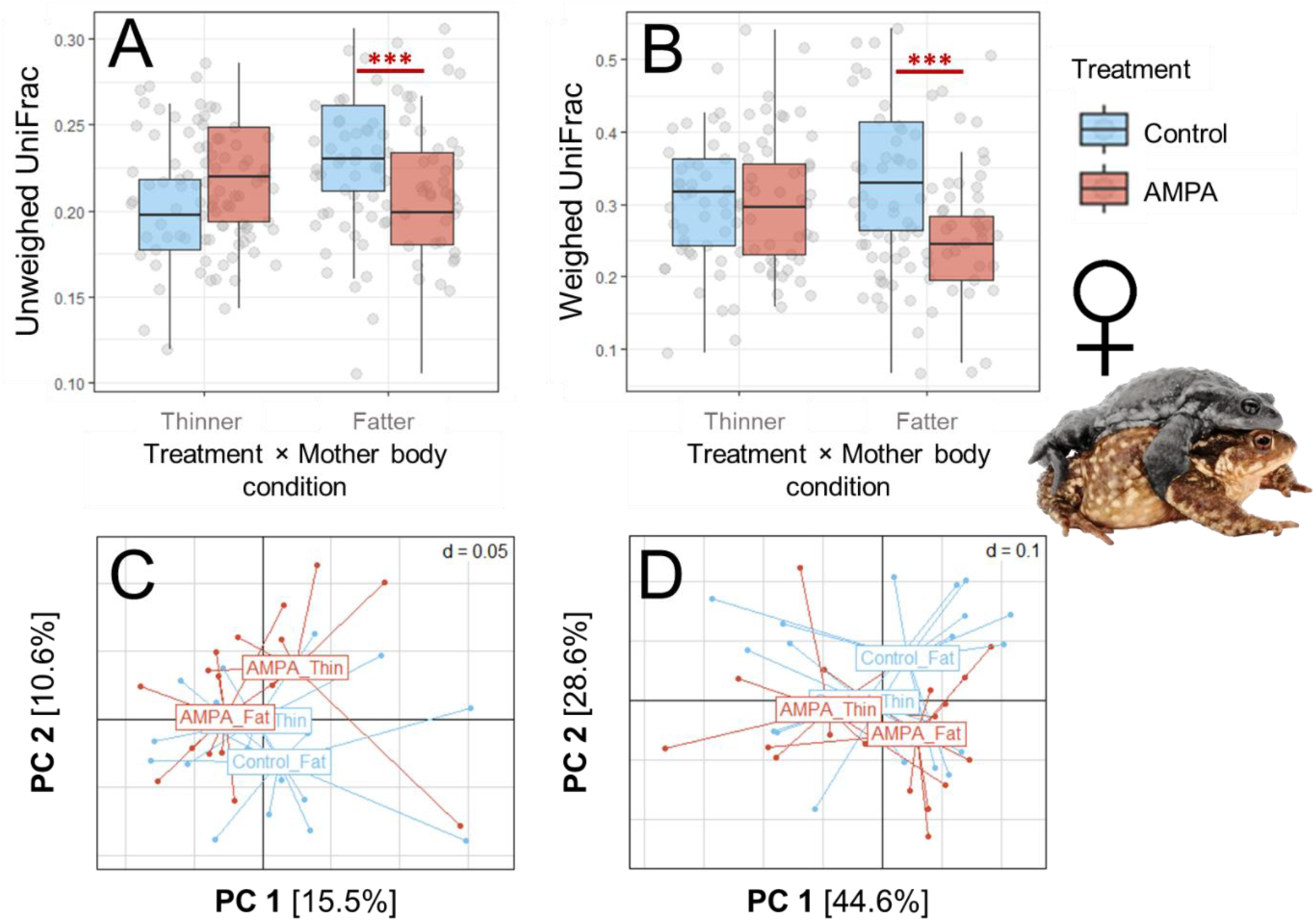
Interaction of treatment and maternal body condition on GS 37 *Bufo spinosus* tadpoles’ faecal microbiota community. The upper panels represent calculated unweighted (A) and weighted (B) UniFrac distances according to the treatment (control in blue vs AMPA in red) and maternal body condition. Lower panels represent principal coordinate analysis (PCoA) for each pool of siblings (AMPA, n = 20, control group, n=20) according to treatment and maternal body condition. Closer dots in the PCoA figure indicate a more similar microbial community. The percentage of variation explained by principal coordinates (PC) is shown on the axes. We calculated body condition from the scaled mass index. Condition categories were separated by median value (’thinner’ ≤ median value; ’fatter’> median value). Significant differences are represented by *** (p<0.007 in A and B; we performed pairwise comparisons using Dunn’s all-pairs test).

**Figure 4:**
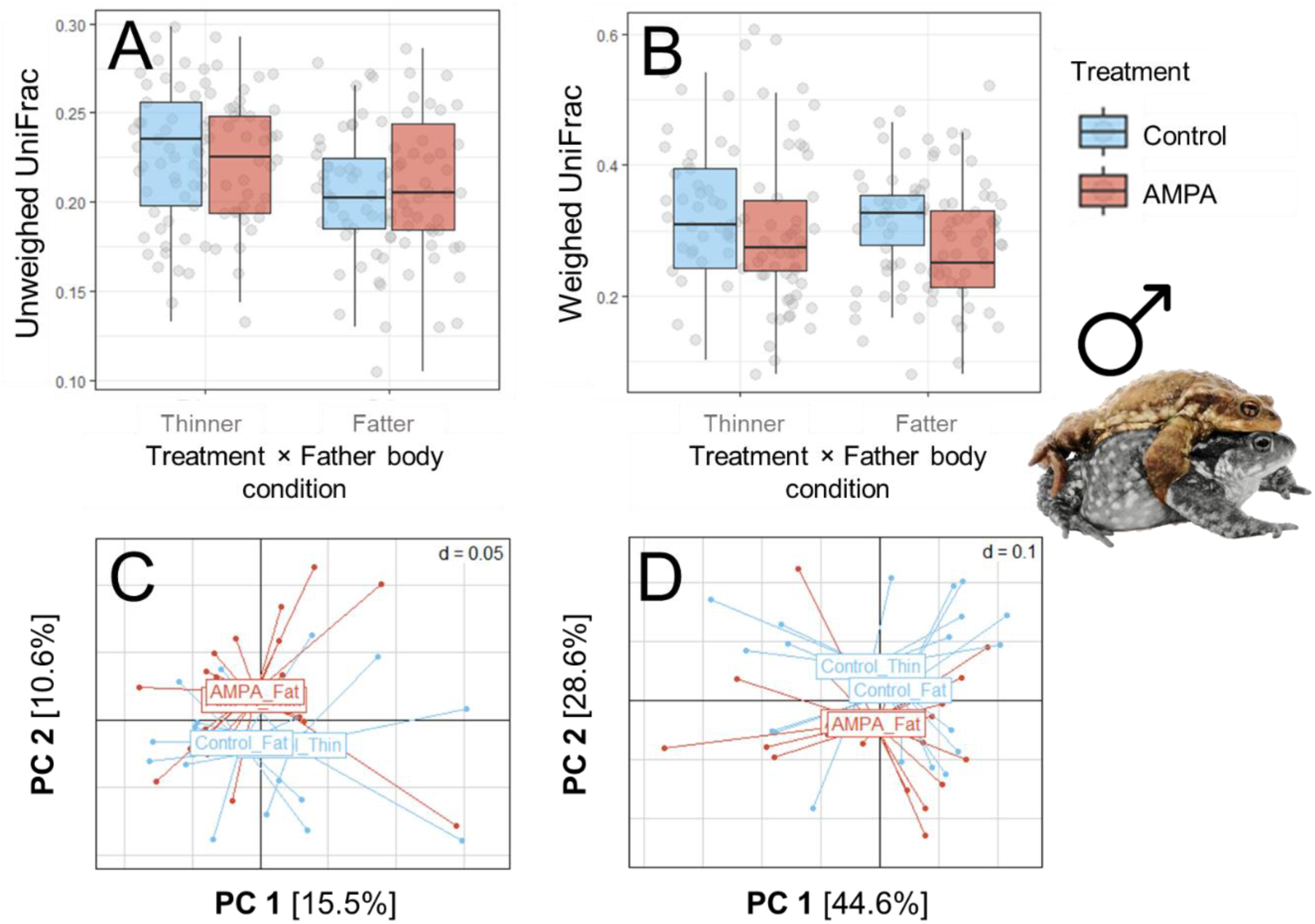
Interaction of treatment and paternal body condition on GS 37 *Bufo spinosus* tadpoles’ faecal microbiota community. The upper panels represent calculated unweighted (A) and weighted (B) UniFrac distances according to the treatment (control in blue vs AMPA in red) and paternal body condition. Lower panels represent principal coordinate analysis (PCoA) for each pool of siblings (AMPA, n = 20, control group, n=20) according to treatment and paternal body condition. Closer dots in the PCoA figure indicate a more similar microbial community. The percentage of variation explained by principal coordinates (PC) is shown on the axes. Body condition was calculated from the scaled mass index. Condition categories were separated by median value (’thinner’ ≤ median value; ’fatter’> median value).

### Interaction between treatment and maternal body condition on tadpole’s microbiome

In AMPA-exposed tadpoles, the abundance of Bacteroidota decreased with increasing maternal body condition (**Figure 5A**, **Table 1**). In contrast, we observed a positive relationship between Bacteroidota abundance and maternal body condition in control tadpoles (**Figure 5A**, **Table 1**).

**Figure 5:**
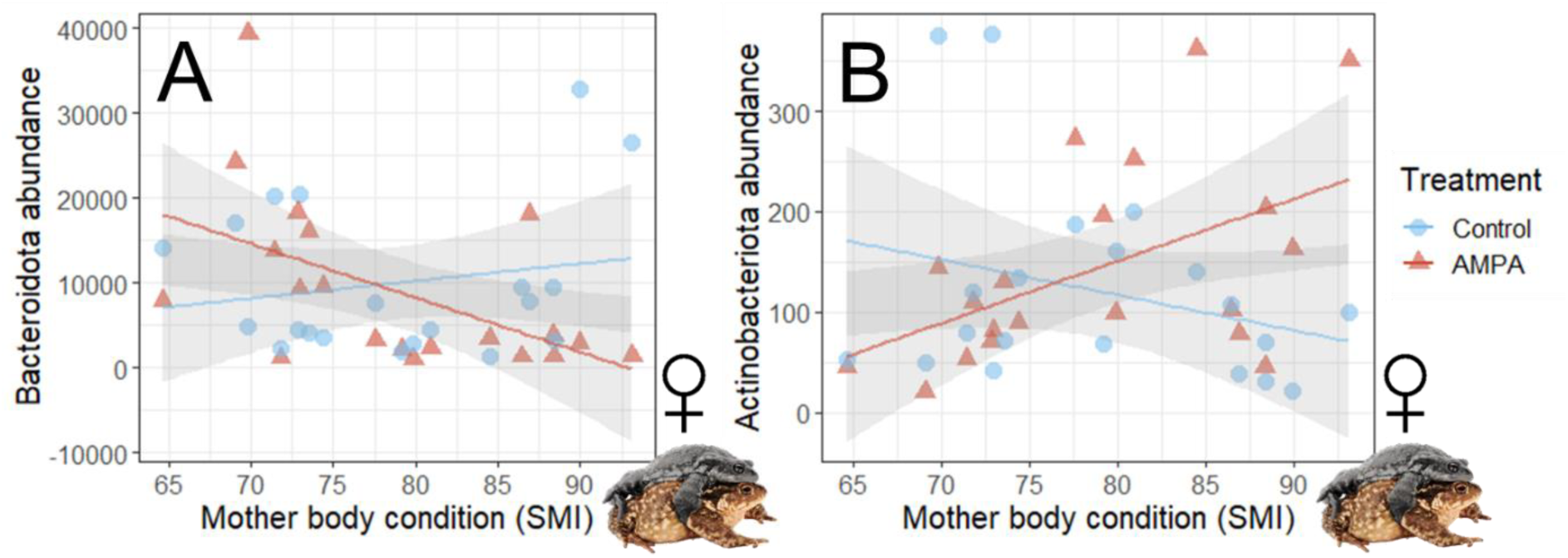
Relationships between tadpoles’ faecal phylum abundance and maternal body condition according to AMPA exposure. Bacteroidota (A) and Actinobacteriota (B) abundances (number of reads) were differently associated with maternal body condition as inferred by their scaled mass index (SMI) according to AMPA exposure (A, conditional r² = 0.23 and B, conditional r² = 0.46). Each red triangle (AMPA exposed) and blue dot (control) represent a pool of faeces obtained from three siblings exposed to a similar treatment.

**Table 1:**
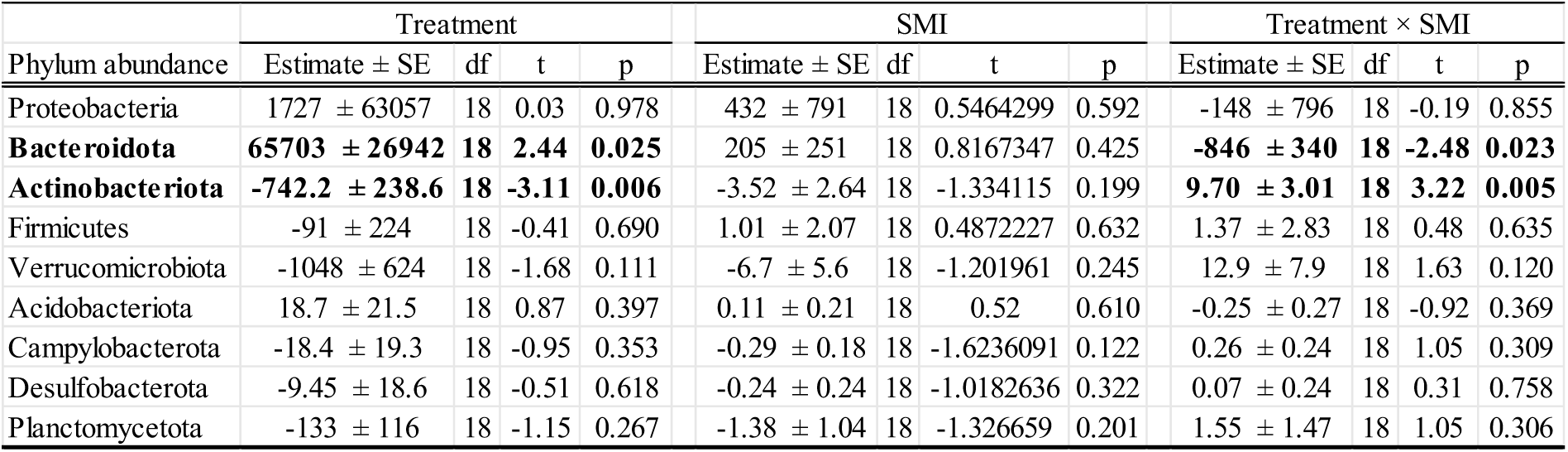
Relationships between the abundance of the nine major phyla sequenced in tadpole’s faecal microbiome, according to AMPA exposure and maternal body condition. Values were obtained using linear mixed effects models with clutch identity as a random factor. Values in bold are significant at the 0.05 level.

In contrast, Actinobacteriota abundance increased with increasing maternal body condition in AMPA-exposed tadpoles, while the Actinobacteriota/maternal body condition relationship was negative in control tadpoles (**Figure 5B**, **Table 1**). The abundance of the seven other phyla was unrelated to treatment, SMI and their interaction (**Table 1**).

The effects of AMPA on microbiota composition were more potent in offspring produced by fatter females (**Figures 3, 5**) within the phyla Bacteroidota and Actinobacteriota (**Table 1**). We, therefore, conducted LEfSe analyses within this subset (i.e. Bacteroidota and Actinobacteriota in the ‘fatter’ female group, **Table 2**). We identified ten markers within the Bacteroidota phylum for the AMPA-exposed group (**Table 2**). The features that explain most the AMPA group were: class Bacteroidia, order Flavobacteriales, family *Weeksellaceae*, genus *Cloacibacterium*, species *cloacibacterium* uncultured, and class Bacteroidia, order Sphingobacteriales, family *KD3-93*, genus *KD3-93 uncultured*. Whereas within the Actinobacteriota phylum, four markers were identified in the AMPA group (**Table 2**), all explained by class Actinobacteria, order Streptomycetales, family *Streptomycetaceae*, genus *Streptomyces* and species *streptomyces*.

**Table 2:** LEfSe analysis on taxonomic biomarkers of gut microbiota influenced by AMPA exposure in offspring from fatter females. LEfSe analysis identified the most differentially abundant taxa within a) Bacteroidota and b) Actinobacteriota according to AMPA exposure. We show LDA scores > 2; k=kingdom, p=phylum, c=class, o=order, f=family, g=genus, s=species, ef. LDA= effect size linear discriminant analysis.

| Marker | Feature | Enriched group | ef. LDA | p-value |
| --- | --- | --- | --- | --- |
| **a) Bacteroidota (Fatter mothers)** |  |  |  |  |
| Marker1 | k__Bacteria\|p__Bacteroidota\|c__Bacteroidia\|o__Flavobacteriales\|f__Weeksellaceae | AMPA | 3.383 | **0.008** |
| Marker2 | k__Bacteria\|p__Bacteroidota\|c__Bacteroidia\|o__Sphingobacteriales\|f__KD3-93 | AMPA | 3.336 | **0.041** |
| Marker3 | k__Bacteria\|p__Bacteroidota\|c__Bacteroidia\|o__Sphingobacteriales\|f__KD3-93\|g__KD3-93_uncultured | AMPA | 3.336 | **0.041** |
| Marker4 | k__Bacteria\|p__Bacteroidota\|c__Bacteroidia\|o__Sphingobacteriales\|f__KD3-93\|g__KD3-93_uncultured\|s__KD3-93_uncultured_uncultured | AMPA | 3.336 | **0.041** |
| Marker5 | k__Bacteria\|p__Bacteroidota\|c__Bacteroidia\|o__Flavobacteriales\|f__Weeksellaceae\|g__Cloacibacterium | AMPA | 3.245 | **0.049** |
| Marker6 | k__Bacteria\|p__Bacteroidota\|c__Bacteroidia\|o__Flavobacteriales\|f__Weeksellaceae\|g__Cloacibacterium\|s__Cloacibacterium_uncultured | AMPA | 3.245 | **0.049** |
| Marker7 | k__Bacteria\|p__Bacteroidota\|c__Bacteroidia\|o__Flavobacteriales\|f__Weeksellaceae\|g__Weeksellaceae_uncultured | AMPA | 2.820 | **0.013** |
| Marker8 | k__Bacteria\|p__Bacteroidota\|c__Bacteroidia\|o__Flavobacteriales\|f__Weeksellaceae\|g__Weeksellaceae_uncultured\|s__Weeksellaceae_uncultured_uncultured | AMPA | 2.820 | **0.013** |
| Marker9 | k__Bacteria\|p__Bacteroidota\|c__Bacteroidia\|o__Flavobacteriales\|f__Flavobacteriaceae\|g__Leptobacterium | AMPA | 2.711 | **0.028** |
| Marker10 | k__Bacteria\|p__Bacteroidota\|c__Bacteroidia\|o__Flavobacteriales\|f__Flavobacteriaceae\|g__Leptobacterium\|s__Leptobacterium | AMPA | 2.711 | **0.028** |
| **b) Actinobacteriota (Fatter mothers)** |  |  |  |  |
| Marker1 | k__Bacteria\|p__Actinobacteriota\|c__Actinobacteria\|o__Streptomycetales\|f__Streptomycetaceae | AMPA | 4.015 | **0.014** |
| Marker2 | k__Bacteria\|p__Actinobacteriota\|c__Actinobacteria\|o__Streptomycetales\|f__Streptomycetaceae\|g__Streptomyces\|s__Streptomyces | AMPA | 4.012 | **0.014** |
| Marker3 | k__Bacteria\|p__Actinobacteriota\|c__Actinobacteria\|o__Streptomycetales | AMPA | 4.012 | **0.014** |
| Marker4 | k__Bacteria\|p__Actinobacteriota\|c__Actinobacteria\|o__Streptomycetales\|f__Streptomycetaceae\|g__Streptomyces | AMPA | 4.009 | **0.014** |

## DISCUSSION

We here show that a minute concentration of AMPA (aminomethylphosphonic acid) - the primary metabolite of glyphosate and the main contaminant detected in surface waters worldwide - can affect gut microbiota biomass and community composition in larvae of a widespread amphibian species. Interestingly, this effect was, at least partly, mediated by maternal condition, as the effects of AMPA on tadpoles’ microbiome were exacerbated in individuals produced by females with better body condition. AMPA did not affect the gut microbiome of tadpoles from leaner females, and paternal body condition was unrelated to the effects of AMPA. The interaction between AMPA and maternal body condition on tadpoles’ gut microbiome was driven by contrasting alterations of the abundance of Bacteroidota and Actinobacteriota, which are respectively the second and the third most abundant phylum in spined toad larval faecal microbiome.

### Possible vertical transmission of the gut microbiota

In amphibian species with aquatic larvae, gut microbiota majorly originates from the environment (i.e. water and diet), and the contribution of parental microbiomes was thought to be minor (Hernández-Gómez and Hua, 2023; Prest et al., 2018; Scalvenzi et al., 2020). In this study, we captured amplectant pairs in four different sites and placed them in a tank filled with dechlorinated tap water until all females laid their eggs. We controlled the tap water, and our analyses indicated that tap water did not contribute to significant amounts of bacterial DNA. We exposed egg strings to the same water and fed tadpoles with the same diet of organic ground spinach. Therefore, the only source of variation in early microbial egg colonizers were those present on the parents (skin and cloacal microbiomes), and we were able to show evidence of a maternal signature on the tadpole faecal microbiome.

Vertical transmission has been extensively studied in mammals, with maternal faecal microbes transferred to newborns during birth (Ferretti et al., 2018; Wampach et al., 2018; Wang et al., 2020). Although prenatal transfer is still debated in oviparous vertebrates, evidence of vertical transmission has been reported. For instance, bacterial colonization during egg formation has been suggested in Eastern Fence Lizard (*Sceloporus undulatus*) (Trevelline et al., 2018), and female *Sceloporus virgatus* lizards transfer beneficial microbes from their cloaca onto their eggs during oviposition with beneficial effects on the offspring (Bunker et al., 2021). Moreover, hatchling loggerhead sea turtles *Caretta caretta* harboured distinct microbial communities with respect to sand and eggshells, suggesting here also a maternal origin of their pioneer gut microbiome (Vecchioni et al., 2022). In addition, the faecal microbiome of neonate Rock pigeons (*Columba livia*) hatched in an incubator resembled the cloacal microbiome of females sampled from the same population (Dietz et al., 2020). Evidence is accumulating from pathogen transmission studies that *Salmonella enterica* contamination of chicken eggs does not occur from penetration through the shell but by the passage from the hen’s intestinal tract to the reproductive tract, then from pathogen colonization into the forming egg on the vitelline membrane, in the egg white or the shell membranes (Gantois et al., 2009). Therefore, maternal intestinal microbiota could colonize the egg yolk before shell deposition.

In contrast to amniotic vertebrates, amphibians produce jelly-coated eggs. Egg-jelly has various functions such as fertilization, insulation, gas exchange, and protection (Altig and McDiarmid, 2007; Beattie, 1980; Burggren, 1985; Olson and Chandler, 1999), yet some pathogenic bacteria can penetrate the thick jelly layer of the egg (Khalifa et al., 2021). Since vertical transmission is possible in shelled eggs, maternal transmission in non-shelled eggs is even more likely, as observed in crustaceans (Giraud et al., 2022). But surprisingly, few researchers have investigated vertical transmission in amphibians without parental care (Hughey et al., 2017). In African clawed frogs (*Xenopus tropicalis)*, the environment was identified as the primary driver of egg bacterial communities by contributing around 70% of the bacteria in controlled conditions. In the same study, the skin and faeces of parents were identified as minor contributors (Scalvenzi et al., 2020). In wild boreal toad populations (*Anaxyrus boreas*), a quarter of the bacterial communities observed on eggs and a third of communities observed on early-stage tadpoles were comprised of bacteria acquired from an unknown source (neither water, upland soil nor sediments), and these strains were likely parentally transmitted (Prest et al., 2018).

Although further studies are needed to understand better the vertical transmission of the gut microbiota in spined toads, substantial evidence points towards this direction. As observed in other taxa, such as mammals and reptiles, vertically transmitted strains are likely to be more ecologically relevant for the offspring compared with non-maternal strains (Ferretti et al., 2018), and transmitted strains could confer fitness advantages to the progeny (Bunker et al., 2021; Trevelline et al., 2018). Mechanisms of transgenerational immune priming could provide complementary explanations (Roth et al., 2018). There is complementary evidence for the transmission of innate immunity compounds (i.e. antimicrobial skin peptides and mutualistic microbiota) from females to eggs in amphibians (Walke et al., 2011). Yet, more studies are needed to investigate and quantify immune responses transmitted to offspring in amphibians.

### Tadpole gut microbiome depends on the maternal body condition

Body condition can be a good indicator of fitness in numerous species (Bowers et al., 2014; Bright Ross et al., 2021; Liu et al., 2020; Milner et al., 2003), including spined toads (Renoirt et al., 2022). Associations between body condition and gut microbiota composition have been well described in human and rodent models, and this is not surprising given the function of the gut microbiome to produce metabolites involved in energy homeostasis and metabolic health (Aron-Wisnewsky et al., 2021.; Fan and Pedersen, 2021; Moreno-Navarrete and Fernandez-Real, 2019; Turnbaugh et al., 2006; Zwartjes et al., 2021); however, in the wild, these relationships are more difficult to characterize.

For instance, no associations between the gut microbiome and body condition were reported in fire salamanders *Salamandra Salamandra*, Seychelles warbler *Acrocephalus sechellensis*, three-spined stickleback *Gasterosteus aculeatus* (Friberg et al., 2019; Wang et al., 2021; Worsley et al., 2021). In contrast, in Eurasian perch *Perca fluviatilis*, lower microbial diversity was related to improved condition; in great tit nestlings *Parus major*, a time-lagged association was observed between gut microbiota composition, nestling weight and survival; in coyotes *Canis latrans,* the consumption of anthropogenic food in urban individuals was associated with increased microbiome diversity, higher abundances of *Streptococcus* and *Enterococcus* and poorer average body condition (Bolnick et al., 2014; Davidson et al., 2021; Sugden et al., 2020), and in wood frogs, *Rana sylvatica* egg microbiome manipulation accelerated larvae growth and development rates (Warne et al., 2019). These contrasted relationships between gut microbiota composition and body condition in wild species could result from environment-dependent variations in feeding activity, diet composition, and body condition, and all these features can also vary according to sex, age, and breeding cycle. Measuring microbiome-fitness relationships at just one point could be misleading in free-ranging species.

In the present study, we observed an association between tadpole faecal microbiome composition and maternal body condition, which was affected by AMPA exposure. This association suggests that components of the maternal microbiomeor determinants of microbiota composition were transmitted to the eggs during oviposition, and this specific microbiome signature was more sensitive to AMPA exposure. We can assume that a gut microbiota composition that is more efficient in harvesting energy, as suggested by the higher maternal body condition in that group, would consist of a gut bacterial assemblage with species more sensitive to AMPA. One may hypothesize that those females in better body condition originate from habitats preserved from AMPA exposition (i.e. forest sites, see Tartu et al. (2022)) and that their gut bacterial composition would be more sensitive to AMPA exposure. However, maternal body condition was not related to habitat (agricultural vs forested, LME estimate: - 2.31 ± 3.69, p = 0.538). The observed relationships underline the dependence of the gut microbiome on interactions among other deterministic factors that were not accounted for in this study, such as host genetic and epigenetic background, age, diet or other environmental stressors (Chen et al., 2022; Shu et al., 2019; Song et al., 2021; Zhou et al., 2021). Nevertheless, we identified a modification of the abundance of two major phyla in tadpole’s gut microbiota, Bacteroidota and Actinobacteriota, which varied according to maternal body condition and AMPA exposure.

### Effects of agrochemicals transformation products according to gut microbiota composition

In line with our hypothesis, AMPA exposure affected tadpoles’ gut microbiome by reducing bacterial biomass and changing community composition. In several taxa,including amphibians, a dysbiosis induced by decreased bacterial biomass is associated with deficient nutrient absorption and impaired immunity (Adamovsky et al., 2018; Gomaa, 2020; Jiménez and Sommer, 2017). In addition, the AMPA – microbiome relationship was exacerbated when transgenerational traits, such as maternal body condition, were considered, highlighting microbial colonizers’ importance in susceptibility to pollutants. Specifically, in tadpoles from better-condition females, we observed a weaker Bacteroidota abundance and a more substantial Actinobacteriota abundance in the AMPA-exposed group.

As previously mentioned, only a few studies conducted on invertebrates have investigated whether AMPA exposure would lead to gut microbiota dysbiosis (Blot et al., 2019; Iori et al., 2020), yet effects of glyphosate exposure similar to those observed in the present study have been reported in Sprague-Dawley rats (Mesnage et al., 2021). For instance, glyphosate exposure decreased Bacteroidota abundance and concomitantly increased Firmicutes and Actinobacteria abundances in rats’ caecum microbiome (Mesnage et al., 2021). The reported effects of glyphosate could also be the consequence of AMPA, as glyphosate is degraded to AMPA in vertebrates and highly accumulates in the intestine, as observed in bird and fish models (Fréville et al., 2022; Yan et al., 2023).

The ability to cleave the C - P bond of AMPA and to use it as a phosphorus source is widespread in bacteria (Dick and Quinn, 1995; Fox and Mendz, 2006; Harkness, 1966; Studnik et al., 2015), including various species of the Actinobacteriota phylumas *Streptomyces* (Obojska et al., 1999; Obojska and Lejczak, 2003). The observed increase of Actinobacteriota is likely to result from the ability of *Streptomyces* and other akin species to utilize AMPA as a phosphorus source, promoting their growth. In contrast, the decreased abundance of Bacteroidota in tadpole gut microbiota, and more specifically the orders Sphingobacteriales (family *KD3-93*) and Flavobacteriales (family *Weeksellaceae*, genus *Cloacibacterium*) could either result from a higher sensitivity of these orders to AMPA, a modification of the gut environment which became less favourable to their growth (e.g. pH) or resource competition with Actinobacteriota (Firrman et al., 2022). This gut microbiota alteration associated with AMPA exposure can have important implications for the host’s health.

For instance, Sphingobacteriales can produce sphingolipids that regulate the immune system and lipid metabolism (An et al., 2011; Bai et al., 2023; Olsen and Jantzen, 2001). Flavobacteriales, on the other hand, play several roles in various metabolic pathways, including vitamins, amino-acid and fatty acid biosynthesis (Rosas-Pérez et al., 2014; Yang et al., 2017; Zhou et al., 2022). Flavobacteriales can thus bear positive effects on the host growth and development (Pan et al., 2023). At the genus level, *Cloacibacterium* sp. can degrade cellulose and may have a critical role in transforming plant-derived complex dietary carbohydrates into essential short-chained fatty acids (SCFA) for herbivoreorganisms such as spined toad tadpoles (Flint et al., 2012; Fujimori, 2021; Hu et al., 2021; Martens et al., 2011; Zhang et al., 2018). Therefore, a decrease in Bacteroidota could disrupt nutrient intakes, leading to a delayed development length, as observed in agricultural AMPA-exposed tadpoles from the present study (Tartu et al., 2022). In the crucian carp (*Carassius auratus*), for instance, glyphosate exposure resulted in a dysbiosis of Bacteroidota at the phylum level, and Bacteroidota abundance was negatively correlated with different metrics of growth performance (condition factor, fat ratio and specific growth rate) (Yan et al., 2022).

We have previously reported that AMPA exposure was associated with a higher deformity rate upon hatching, especially in individuals from AMPA-free forest habitats, and increased development length in AMPA-exposed individuals from agricultural sites (Tartu et al., 2022). While embryonic stages may be more sensitive to AMPA exposure in forest individuals (AMPA-preserved population), they might be more resilient to a gut microbiome dysbiosis as no further effects on fitness were observed at metamorphosis (Tartu et al., 2022). In contrast, agricultural individuals (AMPA-exposed population) could be more resistant during embryonic stages. However, gut microbiome dysbiosis could still result in a longer development duration (Tartu et al., 2022). These findings again underline the part that may play the host genotype in shaping the consequences of gut microbiota dysbiosis. Yet, we have to remain cautious as we only followed the exposed individuals until metamorphosis and deleterious effects could appear later in life, as early-life microbiota composition shapes fitness trajectories in amphibians (Knutie et al., 2017; Warne et al., 2019). In addition, there is alarming evidence of the disappearance of breeding spined toads in agricultural habitats (Renoirt et al., 2024), which could be a long-term effect of early-life exposure to toxicants.

## ACKNOWLEDGEMENT

The authors thank Matthias Renoirt and Marion Cheron for their help on the field and tadpole monitoring, Léa Lorrain-Soligon for providing a *Bufo spinosus* picture, three anonymous referees and Lauris Evariste for their constructive comments that greatly improved previous versions of the manuscript.

## FUNDING

Funding was provided by the CNRS, the Agence de l’Eau Loire-Bretagne, the Agence de l’Eau Adour-Garonne, the Région Nouvelle-Aquitaine (Aquastress 2018-1R20214, Amphitox 2019-1R20216), the ANSES (BiodiTox project # 2019/1/031), The Plan d’Action National ECOPHYTO (n°OFB-21-0941).

## AUTHOR CONTRIBUTION

F.B. obtained the funding and supervised the study; F.B., N.P. and S.T. designed the study; S.T., G. B. and I.C. generated the libraries; N.P. performed bioinformatic analyses; S.T. collected, analyzed, interpreted the data, wrote the first version of the manuscript. All authors have revised and approved the submitted version of the manuscript.

## CONFLICT OF INTEREST

The authors have no conflict of interest to declare.

## ETHICS STATEMENT

We followed all applicable institutional and national guidelines for the care and use of animals. This work was approved by the French authorities (COMETHEA ethic committee and Ministère de L’Enseignement Superieur, de la Recherche et de L’innovation) under permits APAFIS#13477–2018032614077834 and DREAL/2020D/8041.

## DATA, SCRIPTS AND SUPPLEMENTARY INFORMATION AVAILABILITY

We posted the dataset and R scripts used for data analysis on Zenodo at https://doi.org/10.5281/zenodo.10401610. Supplementary information describing the fieldwork sampling design is available with the manuscript.

## Notes

https://www.ebi.ac.uk/ena/data/view/PRJEB71117

https://doi.org/10.5281/zenodo.10401610

### Competing Interest Statement

The authors have declared no competing interest.

### Summary of Updates

This version has been recommended by PCI, we added the PCI badge on the title page.

