## Supplementary material for "Maternal body condition affects the response of larval spined toads’ faecal microbiome to a widespread contaminant": Figure S1; Figure S2

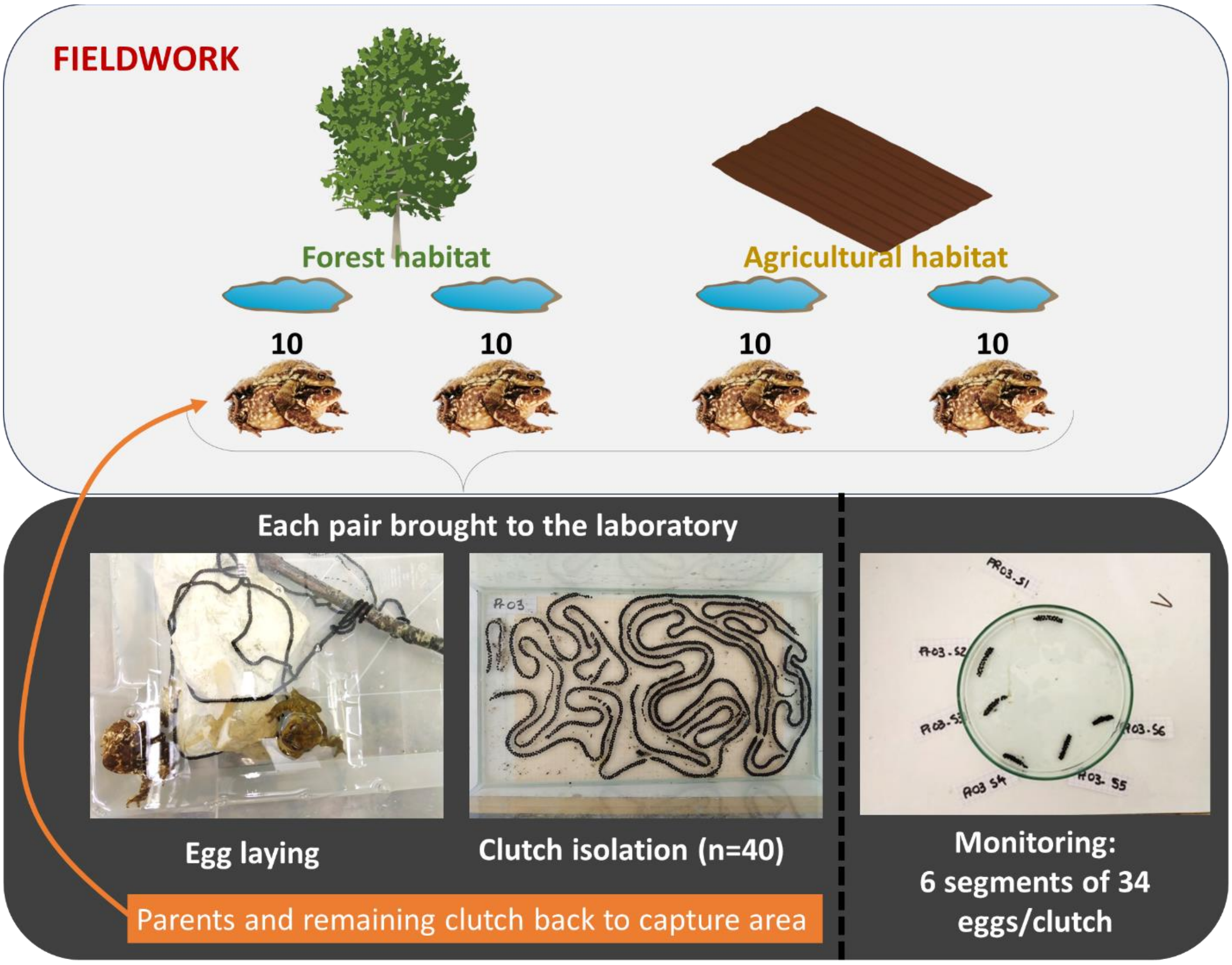


**Figure S1**: Fieldwork design. Forty amplectant pairs were captured in four different sites (two ponds in forested areas and two ponds in agricultural areas) and brought back to the laboratory until the female laid. Clutches were then isolated and six segments of 34 eggs were selected for the AMPA exposure study.

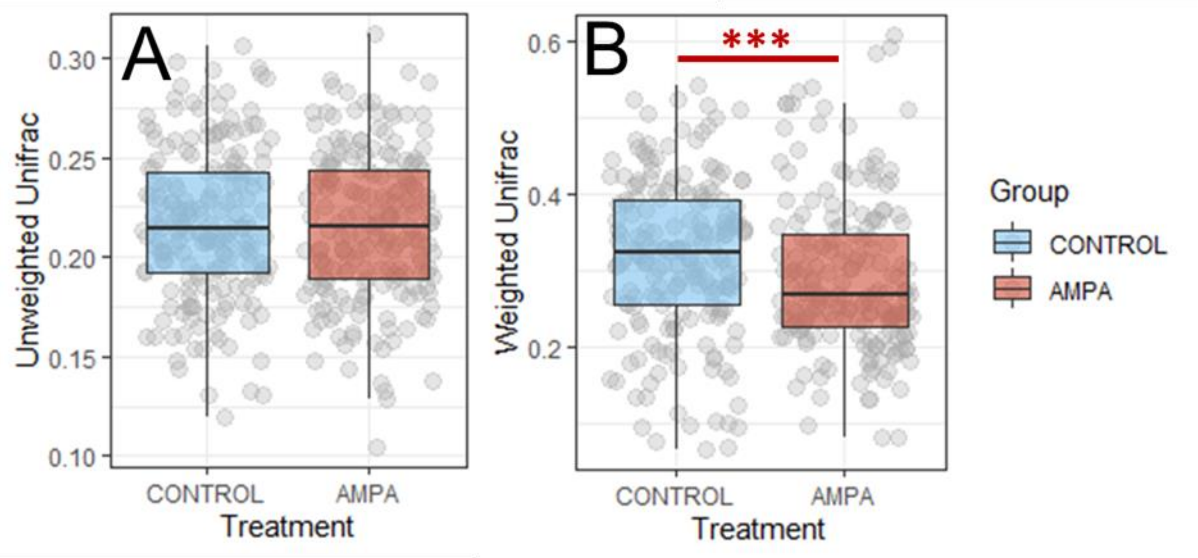


**Figure S2: Relationships between treatment and GS 37 *Bufo spinosus* tadpoles' faecal microbiota community.** Represented data are calculated unweighted (A) and weighted (B) UniFrac distances according to the treatment (control in blue vs AMPA in red). Significant differences are represented by *** (p=0.003 in B).
